# The GluA1 cytoplasmic tail regulates intracellular AMPA receptor trafficking and synaptic transmission onto dentate gyrus GABAergic interneurons, gating response to novelty

**DOI:** 10.1101/2024.12.01.626277

**Authors:** Gerardo Leana-Sandoval, Ananth V. Kolli, Carlene A. Chinn, Alexis Madrid, Iris Lo, Matthew A. Sandoval, Vanessa Alizo Vera, Jeffrey Simms, Marcelo A. Wood, Javier Diaz-Alonso

**Affiliations:** Department of Anatomy & Neurobiology, University of California at Irvine, CA, 92697, USA; Center for the Neurobiology of Learning and Memory, University of California at Irvine, CA, USA; Department of Neurobiology & Behavior, University of California at Irvine, CA, 92697, USA; Gladstone Institute of Neurological Disease, San Francisco, CA 94158, USA

**Keywords:** AMPA receptor, GluA1, C-tail, Carboxy-terminal domain, schizophrenia, dentate gyrus, novelty response, LTP, intracellular trafficking, PV+ interneuron

## Abstract

The GluA1 subunit, encoded by the putative schizophrenia-associated gene GRIA1, is required for activity-regulated AMPA receptor (AMPAR) trafficking, and plays a key role in cognitive and affective function. The cytoplasmic, carboxy-terminal domain (CTD) is the most divergent region across AMPAR subunits. The GluA1 CTD has received considerable attention for its role during long-term potentiation (LTP) at CA1 pyramidal neuron synapses. However, its function at other synapses and, more broadly, its contribution to different GluA1-dependent processes, is poorly understood. Here, we used mice with a constitutive truncation of the GluA1 CTD to dissect its role regulating AMPAR localization and function as well as its contribution to cognitive and affective processes. We found that GluA1 CTD truncation affected AMPAR subunit levels and intracellular trafficking. ΔCTD GluA1 mice exhibited no memory deficits, but presented exacerbated novelty-induced hyperlocomotion and dentate gyrus granule cell (DG GC) hyperactivity, among other behavioral alterations. Mechanistically, we found that AMPAR EPSCs onto DG GABAergic interneurons were significantly reduced, presumably underlying, at least in part, the observed changes in neuronal activity and behavior. In summary, this study dissociates CTD-dependent from CTD-independent GluA1 functions, unveiling the GluA1 CTD as a crucial hub regulating AMPAR function in a cell type-specific manner.

## Introduction

AMPA receptors (AMPAR) mediate moment-to-moment excitatory synaptic transmission at synapses throughout the CNS. Additionally, specific and sustained increases in the postsynaptic AMPAR complement underlie long-term potentiation (LTP) (Kauer, Malenka et al. 1988, Muller, Joly et al. 1988), which plays a crucial role in forms of learning and memory (Martin, Grimwood et al. 2000, Nicoll 2017, Gall, Le et al. 2024). AMPARs assemble into heterotetramers of pore-forming subunits (GluA1-4), decorated by auxiliary subunits. Subunit composition imparts AMPARs’ biophysical properties and trafficking behavior (Malinow and Malenka 2002, Collingridge, Isaac et al. 2004, Diering and Huganir 2018, Hansen, Wollmuth et al. 2021, Bessa-Neto and Choquet 2023). At hippocampal CA1 synapses, GluA1-containing AMPAR are crucial for activity-dependent synaptic trafficking and LTP (Zamanillo, Sprengel et al. 1999, Hayashi, Shi et al. 2000, Shi, Hayashi et al. 2001). However, AMPAR subunit composition varies dramatically among cell types and brain regions (Schwenk, Baehrens et al. 2014), and our understanding of the mechanisms underlying AMPAR trafficking and function at other synapses, particularly at synapses onto inhibitory neurons, is limited.

Structurally, AMPAR subunits contain an amino-terminal domain (ATD, a.k.a. NTD), a ligand-binding domain (LBD), a transmembrane domain which forms the pore channel, and a carboxyl-terminal domain (CTD). Of all these regions, the CTD is the most sequence-diverse, and has therefore received considerable attention by researchers studying subunit-specific AMPAR trafficking rules (Malinow and Malenka 2002, Diering and Huganir 2018, Diaz-Alonso and Nicoll 2021, Bessa-Neto and Choquet 2023, Stockwell, Watson et al. 2024). The GluA2 CTD plays an important role in synaptic scaling (Gainey, Hurvitz-Wolff et al. 2009, Ancona Esselmann, Diaz-Alonso et al. 2017), and the GluA4 CTD regulates its subcellular distribution (Boehm, Kang et al. 2006, Luchkina, Coleman et al. 2017). However, it is the GluA1 CTD which has received most of the attention. GluA1 CTD interactions with Protein 4.1N and Sap97 can regulate intracellular AMPAR trafficking and synaptic content (Shen, Liang et al. 2000, Sans, Racca et al. 2001, Kay, Tsan et al. 2022, Bonnet, Charpentier et al. 2023). During LTP, the GluA1 CTD undergoes phosphorylation by CaMKII, PKC and PKA (Barria 1997, Hayashi 2000, Esteban, Shi et al. 2003), and double phospho-null mutation of Serine 831 and 845 in the GluA1 CTD has been shown to block LTP (Lee, Takamiya et al. 2003). These and other studies support an essential role for the GluA1 CTD in LTP. However, other evidence suggests a more nuanced role: i) the discovery that CTD (Ser 831 / Ser 845)-phosphorylated GluA1 accounts for a negligible fraction of GluA1 at synapses *in vivo* (Hosokawa, Mitsushima et al. 2015) [although another study reported a sizable proportion of phosphorylated GluA1 (Diering, Heo et al. 2016)], ii) the finding that GluA1 lacking the PDZ-binding motif traffics normally (Kim, Takamiya et al. 2005, Kerr and Blanpied 2012). iii), the demonstration that CTD-lacking GluA1 can support basal AMPAR transmission and LTP at hippocampal CA3➔CA1 synapses (Granger, Shi et al. 2013, Diaz-Alonso, Morishita et al. 2020, Watson, Pinggera et al. 2021). Altogether, the emerging picture is that the presence of the GluA1 CTD is unlikely to be an absolute requirement for AMPAR-mediated synaptic transmission and LTP at CA1 PNs, where it may instead play a more subtle role (Diaz-Alonso and Nicoll 2021, Bessa-Neto and Choquet 2023, Stockwell, Watson et al. 2024). However, the contribution of the GluA1 CTD to synaptic transmission at other synapses, especially excitatory synapses onto inhibitory neurons, remains largely unexplored.

The link between glutamatergic dysfunction and neuropsychiatric disorders is well-established (Coyle 2006, Lisman, Coyle et al. 2008, Tamminga, Southcott et al. 2012). Specifically, the GRIA1 gene, which encodes the GluA1 subunit, has been identified as a risk locus for schizophrenia in genome-wide association studies (Ripke, O’Dushlaine et al. 2013, Schizophrenia Working Group of the Psychiatric Genomics 2014), and postmortem analyses of individuals with schizophrenia show reduced levels of GluA1 in several brain regions, including the hippocampus (Harrison 1991, Eastwood 1996, Yonezawa, Tani et al. 2022). Excitatory synaptic plasticity, most importantly LTP, is disrupted in CA1 in GluA1 KO mice, which also exhibit alterations in novelty and salience processing and working memory reminiscent of some of the symptoms of schizoaffective disorders (Zamanillo D.; Sprengel and Kaiser 1999, Reisel, Bannerman et al. 2002, Bannerman, Deacon et al. 2004, Sanderson, Sprengel et al. 2011, Barkus, Feyder et al. 2012, Barkus, Sanderson et al. 2014, Bannerman, Borchardt et al. 2018, Panayi, Boerner et al. 2023).

Using GluA1 CTD truncated (ΔCTD GluA1) mice, we found that the GluA1 CTD regulates AMPAR subunit protein levels and subcellular distribution. Interestingly, the CTD is required for some GluA1-dependent functions, most notably the regulation of the response to novelty as well as anxiety- and despair-related behaviors, but not for GluA1-dependent memory processes. Our results suggest that the GluA1 CTD modulates AMPAR synaptic transmission in a subunit composition-dependent and cell type-specific manner. Altogether, this study expands our understanding of the cell-type specific regulation of excitatory synaptic transmission and sheds light into the neurobiological mechanisms regulating the putative schizophrenia risk-associated GluA1.

## Materials and Methods

### Animals

All animal procedures were approved by the Institutional Animal Care and Use Committee at the University of California, Irvine (protocol numbers AUP-20-156; AUP-23-076). Mice were maintained in a 12-hour light/dark schedule and had access to food and water, ad libitum. Generation of homozygous HA-ΔCTD GluA1 knock-in (referred to as ΔCTD GluA1) mice was previously described (Diaz-Alonso, Morishita et al. 2020). Genotyping was carried out by TransnetYX Inc.

### Biochemistry

WT and ΔCTD GluA1 mouse forebrains were dissected and homogenized in Synaptic Protein Extraction Reagent (SynPER, Thermo Scientific, #87793) with protease inhibitors (cOmplete, Roche, #11836170001). Synaptosomes were then obtained following manufacturer’s instructions, as in (Bernard, Exposito-Alonso et al. 2022). For immunoblot, whole brain lysates and synaptosomal fractions were denatured at 95 °C for 5 min. in Laemmli sample buffer (Sigma, #S-3401) and processed for SDS-PAGE. Immuno-Blot PVDF membranes (Bio-Rad, #1620177) were blocked with 5% blotting grade nonfat milk (Lab Scientific, #M0841) in tris-buffered saline with 0.1% tween 20 (Sigma-Aldrich, #P1379). The following primary antibodies were used at a 1:1000 dilution: guinea pig anti-GluA2 CTD (Synaptic Systems, #182 105), mouse anti-GluA1 ATD (Cell Signaling, #13185S), rabbit anti-GluA3 (Alomone Labs, #AGC-010), rabbit anti-GluA4 (Cell Signaling, # 8070), mouse anti PSD-95 (Synaptic systems, #124 011) and mouse anti-tubulin (Millipore-Sigma, #T9026). HRP-conjugated secondary antibodies raised against the appropriate species were used: anti-rabbit IgG (Vector laboratories #PI-1000), anti-mouse IgG (Vector laboratories #PI-2000), and anti-guinea pig IgG (Millipore Sigma #AP108P). Membranes were incubated with ClarityTM Western ECL (BioRad, #170-5060). When needed, membranes were incubated in stripping buffer containing Guanidine HCl and β-mercaptoethanol and triton x-100 in pH 7.5 Tris HCl buffer, with gentle agitation at RT for 30 min. Following incubation, membranes were rinsed, blocked and incubated with another Ab.

### Confocal microscopy and image analysis

WT and ΔCTD GluA1 brain samples were sectioned (40 µm, coronal) following fixation in 4% paraformaldehyde. After blocking with 5% swine serum (Jackson Immuno Research, #014-000-121) and 2% BSA (Cell Signaling, #9998S) in permeabilizing conditions (0.1% Triton X-100, Sigma-Aldrich, #T8787), samples were incubated overnight at 4° C with the following primary antibodies: rabbit anti-GluA1 ATD (Cell signaling, #13185, 1:500), guinea pig anti-GluA2 (Synaptic Systems, #182 105, 1:500), rabbit anti-GluA3 (Alomone Labs, #AGC-010, 1:500), rabbit anti-GluA4 (Cell Signaling, #8070, 1:500), rabbit anti c-Fos (Abcam, #AB190289, 1:500) and mouse anti PSD-95 (Synaptic Systems, #124 011, 1:500) followed by incubation with Alexa 488 goat anti-mouse (Life Technologies, #A-11001, 1:500), Alexa 594 goat anti-rabbit (Life Technologies, #A11012, 1:500), Alexa 647 goat anti-rabbit (Life Technologies, #A21245, 1:500) and Alexa 568 goat anti-guinea pig (Life Technologies, #A11075, 1:500) secondary antibodies for 2 hours at RT. Slides were mounted with ProLong Gold Antifade Reagent with DAPI (Cell Signaling Technology, #8961S).

Confocal images were collected using a Leica Sp8 confocal microscope (Leica Microsystems, Wetzlar, Germany). Dorsal hippocampus field CA1 images including stratum pyramidale and stratum radiatum (SR) were acquired using a 63x oil objective as a series of 20 z-steps, with a z-step size of 1 μm, at a resolution of 1024 x 1024 pixels, and a scanning frequency of 400 Hz. The optical resolution (voxel size) per image was 180 nm in the xy-plane and 1.03 μm in the z-plane. Analysis of synaptic localization was performed using Imaris 9.9.1 (Bitplane, South Windsor, CT, USA) and MatLab Runtime R2022b (Mathworks, Natick, MA, USA), as previously described (Bemben, Sandoval et al. 2023). Briefly, the “Spots” tool was utilized to assign representative three-dimensional ellipsoid shapes to individual synaptic-like GluA1, GluA2, GluA3 and PSD-95 puncta. Then “Background Subtraction” was applied to reduce background signal. A region of interest (ROI) was created to restrict the colocalization quantification to CA1 SR. The number of spots was adjusted qualitatively using the automatically generated and interactive “Quality” filter histogram to select dense signal while excluding puncta likely to be background signal. To ensure an accurate spot segmentation of the underlying puncta determined by size, the “Different Spots Sizes” selection was utilized, adjusting contrast with the “Local Contrast” tool. The histogram was adjusted to accurate puncta coverage. Spots were then rendered. Once optimal settings for each of these parameters were established for the GluA1, GluA2, GluA3, or PSD-95 channels, a batched protocol to automate spot detection on every image was run. Threshold for colocalization was established at 0.7 μm from the center of neighboring puncta.

### Electrophysiology

Whole-cell patch-clamp recordings were obtained from DG GCs or GABAergic interneurons (INs) using acute brain slices from 2-6 months-old male and female mice. 300 µm horizontal slices were obtained in ice-cold, oxygenated NMDG recovery solution containing (in mM): 92 NMDG, 2.5 KCl, 1.25 NaH_2_PO_4_, 30 NaHCO_3_, 20 HEPES, 25 glucose, 2 thiourea, 5 Na-ascorbate, 3 Na-pyruvate, 0.5 CaCl_2_•2 H_2_O, and 10 MgSO_4_•7 H_2_O. pH was adjusted to 7.4 and osmolarity to 310-316 mOsm. Slices were then incubated for at least 30 min. at 34°C in artificial cerebrospinal fluid (aCSF) composed of (in mM): 119 NaCl, 2.5 KCl, 1 NaH_2_PO_4_, 26.2 NaHCO_3_, 11 glucose, 2.5 and 1.3 MgSO_4_. aCSF was bubbled with 95% O_2_ and 5% CO_2_. Osmolarity was adjusted to 307-310 mOsm. For recordings, slices were perfused with aCSF containing 100 µM picrotoxin to block GABA A-mediated responses. Recording pipettes (3-6 MΩ) were filled with internal solution containing (in mM): 135 CsMeSO_4_, 8 NaCl, 10 HEPES, 0.3 EGTA, 5 QX-314, 4 Mg-ATP, 0.3 Na-GTP, and 0.1 spermine. Osmolarity was adjusted to 290-292 mOsm, and pH at 7.3–7.4. Membrane holding current, input resistance and pipette series resistance were monitored throughout experiments. Data were gathered through a IPA2 amplifier/digitizer (Sutter Instruments), filtered at 5 kHz, and digitized at 10 kHz. Series compensation was not performed during data acquisition. For evoked EPSC recordings, a tungsten bipolar electrode was placed in the DG stratum moleculare (SM), thereby stimulating perforant path (PP) inputs onto DG GCs. Electric pulses were delivered at 0.2 Hz. AMPAR EPSCs were obtained while holding the cell at −70 mV; NMDAR currents were obtained at +40 mV. The peak evoked AMPAR response and NMDAR component 100 ms after the stimulation artifact (to avoid contribution of the AMPAR EPSC) were used to calculate the AMPAR/NMDAR ratio. In paired-pulse ratio (PPR) experiments, stimulation was delivered at an inter-stimulus interval of 50 ms. PPR was calculated by dividing the second EPSC by the first. Input/Output (I/O) relationship was assessed by stimulating PP in increments of 50 µA, from 0 µA to 500 µA. For long-term potentiation (LTP) experiments, after obtaining a stable baseline, LTP was induced, no more than 6 min. after break-in, using a theta-burst stimulation (TBS) induction protocol, consisting in four trains of TBS, each train comprised of 5 bursts of spikes (4 pulses at 100 Hz) at 5 Hz applied to the SC fibers at 0.1 Hz, paired with postsynaptic depolarization at 0mV, as in (Traunmuller, Gomez et al. 2016). Statistical analysis was performed at min. 45 after induction. Recordings from cells lost at any point between induction and the end of the experiment (min. 40) were considered until that point.

Electrophysiology data was gathered and analyzed using Sutterpatch (Sutter Instruments) and Igor Pro (Wavemetrics).

### Behavior

Mice were group-housed with littermates. Mice were handled for 1 min for 4 consecutive days prior to all behavioral testing. At the beginning of each testing day, mice were allowed to acclimate to the behavior room for at least 30 min. before the start of the experiment. Behavioral chambers and objects were cleaned and de-odorized between mice. Behavioral scoring was done by a researcher blind to the genotype. Initial behavioral assessments performed at the Gladstone Institute Behavior Core used male mice only. Subsequent studies at UC Irvine included both male and female mice, and data from both sexes were pooled.

#### Open Field (OF)

Mice were placed at the center of an OF arena and allowed to explore for 15 min. In the Gladstone experiments, activity was recorded in a clear acrylic (41 x 41 x 30 cm) chamber using a Flex-Field/Open Field Photobeam Activity System (San Diego Instruments, San Diego, CA) with two 16 x 16 photobeam arrays that automatically detected horizontal and vertical (rearing) movements. Rearings were also quantified. In the UCI experiments, locomotor activity was recorded by an overhead camera in a white, 30 x 23 x 23 cm plastic chamber and total distance traveled was analyzed using a tracking analysis code written in MatLab (Github: https://github.com/HanLab-OSU/MouseActivity). The center / total movement ratio was calculated.

#### Object Location Memory (OLM) task

Mice were habituated to a white Plexiglas chamber (30 x 23 x 23 x cm) for 5 min. daily for 4 days. On the training day, mice were placed in the chamber with two identical objects and allowed to explore them for 10 min. On the test day, 24 hours later, mice were placed in the chamber with either object displaced to a different location and allowed to explore the arena for 5 min. Object identity was counterbalanced between genotypes. The animal’s behavior was recorded using an overhead camera and object exploration time scored using the criteria described by (Vogel-Ciernia and Wood 2014). Discrimination index (DI) was calculated as follows: (Novel Object Time – Familiar Object Time) / (Novel Object Time + Familiar Object Time) x 100. A DI score of +20 or greater was determined as learning. DI was calculated for both training and test day. Exclusion criteria: Mice that scored ±20 preference for an individual object on training day and mice that explored the objects less than 3 seconds were excluded from analysis.

#### Novel Objection Recognition (NOR) task

Mice handling and habituation were as described for the OLM task. On training day, mice were placed in the chamber with two identical objects and allowed to explore them for 10 min. The following day (test day), mice were placed back in the chamber with one familiar and one novel object and allowed to explore for 5 min. The identity of the novel object was counterbalanced between genotypes. Discrimination index was calculated as described for OLM.

#### Forced Alternation Y-maze

The forced alternation task was performed using an opaque Plexiglas Y-maze. Each arm was 36 x 21 x 10 cm. On the training trial, mice were placed into a starting arm, facing the center of the maze, and allowed to explore two of the arms for 5 min., while the third arm was blocked. After an inter-trial interval of 1 min., mice were placed back in the maze at the same starting arm and allowed to explore all three arms for 5 min. The starting arm and blocked arm were counterbalanced across mice. The maze was cleaned and deodorized with 70% ethanol between trials. Total number of arm crossings and time spent in each arm was scored using a mouse tracking software (Any-Maze, Stoelting Co). Mice were required to enter an arm with at least 2/3 of its body to be considered a crossing. DI was calculated as Novel Arm Time / (Novel Arm Time + Non-Starting Arm) x 100 (Wolf et al., 2016).

#### Elevated Plus Maze

Mice were placed in the center of an elevated maze with two open arms (without walls, 38 x 5 cm) and two closed arms (with 16.5 cm tall walls), the intersection of the arms was 5 x 5 cm, and the entire maze is elevated 77.5 cm above the ground (Hamilton-Kinder, Poway, CA). Total time spent and distance traveled in each arm were measured across the 10-min session.

#### Forced Swim Test

Mice were individually placed in a clear plastic cylinder (25.5 cm diameter x 23 cm height), filled with water at 24°C, for 5 min. The total time spent immobile in the last 3 min. of the task was scored. Floating, balancing and idle swimming were considered immobility (Can, Dao et al. 2012).

#### Light/Dark Transition Test

The light-dark apparatus consisted of an opaque acrylic box (42 x 21 x 25 cm) divided into two compartments (2/3 light, 1/3 dark) with a small opening connecting the two chambers. The light compartment was made of opaque white walls and lit by an overhead lamp, while the dark compartment was unlit and made of black non-transparent acrylic walls. Mice were first placed in the light compartment and allowed to freely explore both chambers for 10 min. The time spent in each chamber, number of crossings, and the latency to enter the dark chamber was recorded using Any-Maze (Stoelting Co.).

#### Contextual Fear Paradigm

Fear conditioning experiments were conducted using a Med Associates VideoFreeze system. The fear conditioning chamber (24 x 30.5 x 21.5 cm) sits inside a sound attenuating shell (63.5 x 75 x 35.5 cm, Med Associates, Fairfax, Vermont). On the training day, mice were placed into a conditioning chamber and four-foot shocks (0.45 mA, 2s) were delivered at min. 5, 7, 9, and 11 of a 13-minute training period. The following day (context recall test), mice were exposed to the conditioned context in the absence of foot shocks for 10 min. Fear generalization was assessed 48 hours after the initial training in a different context in a 10 min. session. In this generalization context, tactile, visual, auditory, and olfactory stimuli were all distinct from the training context. Freezing behavior was measured at baseline and during conditioning, the contextual recall test, and the generalization test.

For the pre-exposure experiment, on the pre-exposure day mice were placed into the conditioning chamber for 30 min., with no foot shocks. 24 hours later, on conditioning day, mice were placed back into a conditioning chamber for 13 min, with four foot shocks (0.6mA, 2s) delivered at min. 5, 7, 9, and 11. 24 hours later, on the third day, mice were placed into the conditioned context in the absence of foot shocks for a context recall test, where freezing was measured across a 10 min period. The chamber context remained the same over all three days.

Shock reactivity was measured during training by the VideoFreeze system and expressed as the max motion index.

#### Hot plate test

Hot plate nociception was measured on a black anodized, aluminum plate (IITC Life Science, Woodland Hills, CA) heated to 52°C. Latency to withdraw one of the hind paws from the plate was measured to the nearest hundredth of a second.

### Stereotaxic Viral Injection

Mice were anesthetized using isoflurane and bilaterally injected using a pulled glass pipette in the hippocampal DG field (AP: −3.39, ML: ±2.50, DV: −3.4, −2.9, −2.4) with 1 µl pAAV-mDlx-GFP-Fishell-1 (83900-AAV1), kindly shared by Dr. Gordon Fishell (Dimidschstein, Chen et al. 2016) and purchased from Addgene.

### Statistical Analysis

Data analysis throughout the study was done blind to the experimental condition when possible. Results shown represent the mean ± SEM. The number of independent experiments or biological samples, and the statistical test employed, are indicated in every case. Statistical analyses were performed using GraphPad Prism 9 and SutterPatch software.

## Results

### Truncation of the GluA1 CTD affects AMPAR levels and subcellular distribution

Here we set out to investigate the influence of the GluA1 CTD in AMPAR trafficking, synapse type-specific synaptic transmission and plasticity, cognitive function, novelty processing and other behaviors using ΔCTD GluA1 mice (Fig. 1A). First, we examined whether GluA1 CTD truncation affects AMPAR subunit levels. We observed that GluA1 levels were significantly reduced in ΔCTD GluA1 forebrain lysates compared to their WT counterparts’, yet no differences were observed in synaptosome-enriched fractions (Fig. 1B, C). These findings suggest that the loss of the CTD reduces GluA1 expression or stability, but does not alter GluA1’s synaptic content. In contrast, GluA2 levels were strongly upregulated in ΔCTD GluA1 samples, both globally and in the synaptic fraction (Fig. 1B, D). GluA3 levels were unaffected (Fig. 1B, E). Finally, we observed a modest, statistically significant increase in GluA4 levels in ΔCTD GluA1, yet only in synaptic fractions (Fig. 1B, F).

**Figure 1.**
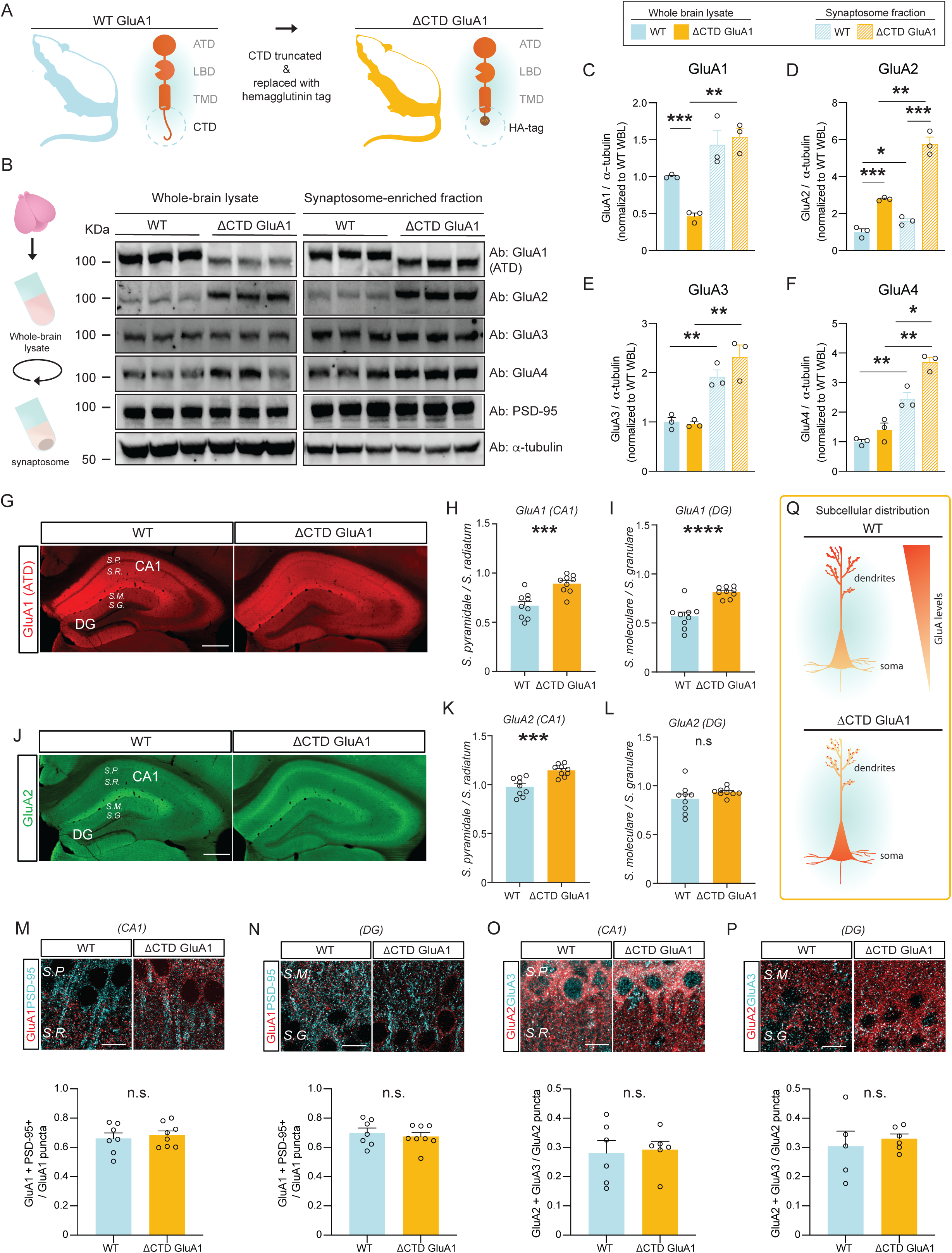
AMPAR subunit levels and subcellular distribution are affected by the loss of the GluA1 CTD. A: Schematic depicting CTD truncation in ΔCTD GluA1 mice. B: Schematic of synaptosomal fractionation (left) and immunoblot from whole-brain lysate (WBL) and synaptosomal fractions of WT and ΔCTD GluA1 (right). C-F: GluA1 (C), GluA2 (D), GluA3 (E), and GluA4 (F) levels normalized to α-tubulin from WT WBL. G: GluA1 ATD staining (red) in WT and ΔCTD GluA1 hippocampus. H-I: Average soma / dendrite ratio of GluA1 signal in CA1 and DG, respectively. J: GluA2 staining (green) in WT and ΔCTD GluA1 hippocampus. K-L: Average soma/dendritic ratio of GluA2 in WT and ΔCTD GluA1 mice for hippocampal field CA1 and DG, respectively. M, N: Representative immunostaining of GluA1 (red) and PSD-95 (cyan) in CA1 and DG in WT and ΔCTD GluA1 samples (top) and colocalization quantification (bottom). O, P: Representative immunostaining of GluA2 (red) and GluA3 (cyan), in CA1 and DG in WT and ΔCTD GluA1 samples (top) and colocalization quantification (bottom). Q: Schematic of subcellular distribution of GluA1 and GluA2 in CA1 and DG in WT and ΔCTD GluA1 PNs. S.P., Stratum pyramidale; S.R., Stratum radiatum; S.M., Stratum moleculare; S.G., Stratum granulare. Scale bar: G, J, 200 µm; M-P, 10 µm. Error bars represent SEM. n.s., not statistically different; *, p≤0.05; **, p≤0.01; ***, p≤0.001; ****, p<0.0001. C-F: one-way ANOVA. H-P: unpaired t-test.

We then examined whether GluA1 CTD truncation affects subcellular AMPAR localization. Using an antibody against the GluA1 ATD, which detects both WT and ΔCTD truncated GluA1, we observed that, as expected, GluA1 immunoreactivity (i.r.) was largely absent from the somata-enriched strata pyramidale (SP) in hippocampal fields CA1-CA3 and granulare (SG) in DG in WT samples. Meanwhile, the subcellular distribution of ΔCTD GluA1 was more diffuse, suggesting impaired intracellular trafficking (Fig. 1G). Quantification of the soma/dendrite GluA1 ir ratio in CA1 and DG revealed a significant accumulation of ΔCTD GluA1 in the soma in both regions (Fig. 1H, I), suggesting that GluA1 CTD truncation impairs AMPAR soma➔dendrite trafficking in CA1 PNs and DG GCs. Interestingly, GluA2 subunits showed a similar redistribution in CA1 (Fig. 1J, K), reminiscent of the pattern found in GluA1 KOs (Zamanillo D.; Sprengel and Kaiser 1999). GluA2 distribution was not significantly altered in DG (Fig. 1J, L). We then turned to confocal microscopy to further analyze GluA1 and GluA2 distribution in field CA1 SR and in DG SM, where most excitatory synapses onto CA1 PNs and DG GCs, respectively, occur. Consistent with our previous observations, we found a significant decrease in the density of putative synaptic GluA1 puncta in both CA1 SR and DG SM (Suppl. Fig. 1A, C). The density of the excitatory postsynaptic marker PSD-95 puncta was slightly reduced in CA1 SR (Suppl. Fig. 1B), but not significantly altered in DG SM (Suppl. Fig. 1D). Despite the significant redistribution of GluA1, its colocalization with PSD-95 was unaffected in both regions in ΔCTD GluA1 samples (Fig. 1M, N), suggesting that ΔCTD GluA1 localization at synapses was not significantly affected. In hippocampal PNs, GluA1/A2 heterotetramers are the most prevalent AMPAR composition, followed by GluA2/A3 (Lu, Shi et al. 2009). To reveal possible compensatory changes in AMPAR subunit composition in ΔCTD GluA1 mice, we assessed the distribution of GluA2 and GluA3. Putative synaptic puncta densities were not altered in CA1 SR or DG SM (Suppl. Fig. 1E-H), and neither was their colocalization (Fig. 1O, P). Altogether, these findings indicate that loss of the GluA1 CTD affects intracellular trafficking, but that the synaptic AMPAR complement is largely intact (Fig. 1Q).

### ΔCTD GluA1 mice exhibit novelty-induced hyperlocomotion but intact cognitive function

Having established the impact of GluA1 CTD truncation in AMPAR levels and subcellular distribution, we sought to clarify whether GluA1-dependent regulation of cognitive function and behavior require the CTD. Previous studies have shown that GluA1 KO mice have impaired spatial working memory, but intact or even enhanced long-term memory (Sanderson, Good et al. 2009). Novelty-induced hyperlocomotion is one of the most robust and reproducible phenotypes in GluA1 KO mice (Zamanillo D.; Sprengel and Kaiser 1999, Bannerman, Deacon et al. 2004, Procaccini, Aitta-aho et al. 2011). To assess the contribution of the GluA1 CTD to spatial novelty processing, we quantified locomotion in the open field (OF) test in WT and ΔCTD GluA1 mice. Initially we tested male WT and homozygous ΔCTD GluA1 mice, and observed a strong exacerbation of novelty-induced locomotion in ΔCTD GluA1 mice compared to WTs (Fig. 2A). The center/total distance ratio was similar in WT and ΔCTD GluA1 mice (Suppl. Fig. 2A). ΔCTD GluA1 mice made significantly fewer fine movements (Suppl. Fig. 2B) and a similar number of rearings (Suppl. Fig. 2C) compared to their WT counterparts. In a different cohort, ΔCTD GluA1 male and female mice showed indistinguishable exacerbated novelty-induced hyperlocomotion, which was absent in heterozygous mice (Suppl. Fig. 2D).

**Figure 2.**
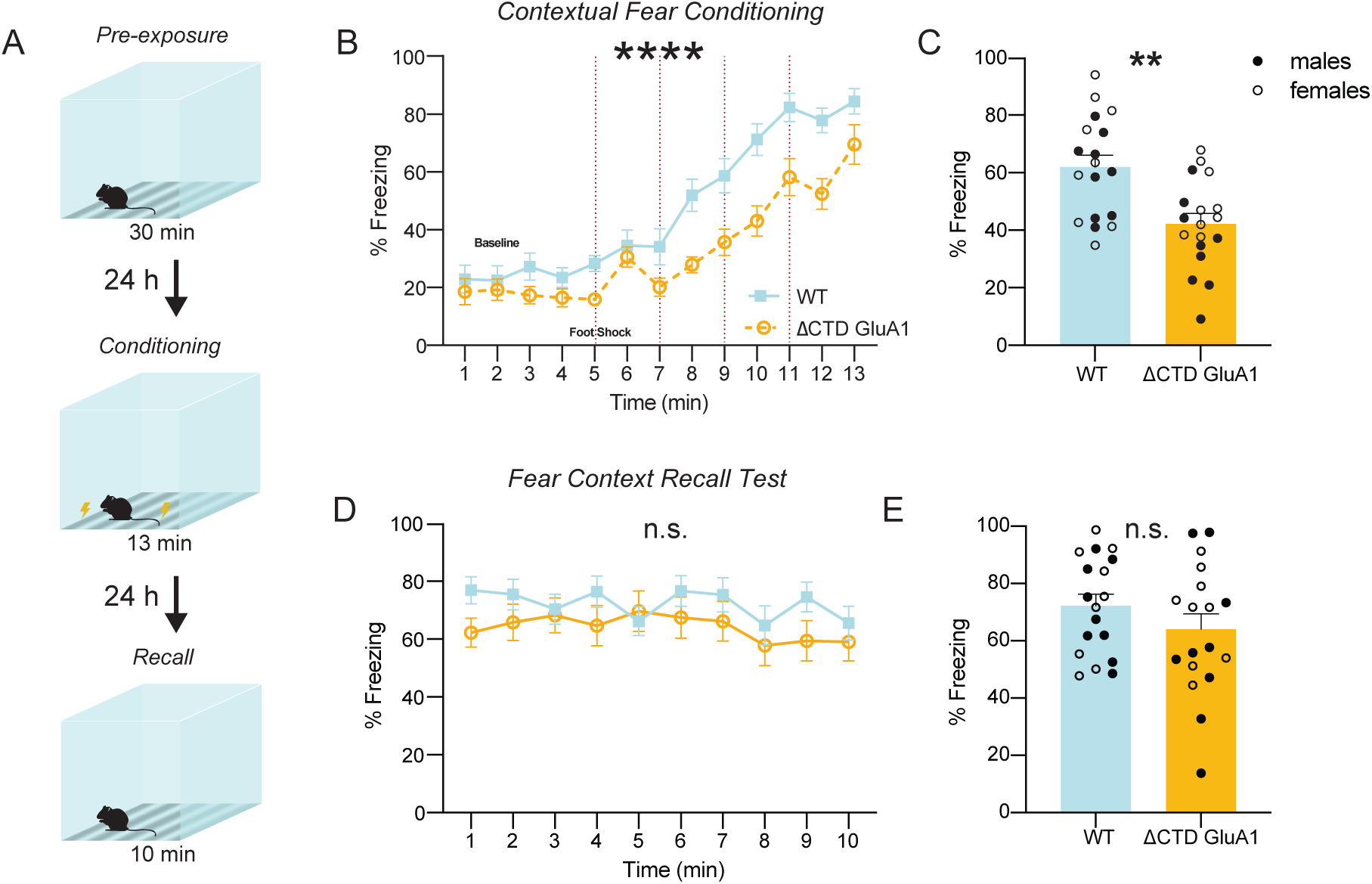
ΔCTD GluA1 mice exhibit novelty-induced hyperlocomotion and impaired fear expression, but intact memory. A: Mean distance traveled during habituation phase for WT and ΔCTD GluA1 mice. B: Schematic of forced alternation Y-maze task. C: Time in novel arm relative to total time n novel and familiar arms for WT and ΔCTD GluA1 mice. D: Representative track plots overlayed atop heat maps of WT (left) and ΔCTD GluA1 (right) mice during habituation day 1 and 4. E: Mean distance traveled across time during habituation for WT and ΔCTD GluA1 mice. F: Schematic of object location memory (OLM) task (left) and representative heat maps (right) of WT and ΔCTD GluA1 mice during training and test day. G: Discrimination index during training and test sessions for WT and ΔCTD GluA1 mice in the OLM task. H: Schematic of novel object recognition (NOR) task (left) and representative heat maps (right) of WT and ΔCTD GluA1 mice during training and test day. I: Discrimination index during training and test sessions for WT and ΔCTD GluA1 mice in the NOR test. J: Schematic of contextual fear conditioning test. K, L: Freezing during training (K) and during the 24-hour contextual recall (L) during contextual fear conditioning for WT and ΔCTD GluA1 mice. Foot shocks are indicated with vertical red dashed lines. M, N: Freezing % across time (M) and average freezing % (N) during context recall test for WT and ΔCTD GluA1 mice. Error bars represent SEM. Empty dots represent females, filled dots represent males. n.s., not statistically different; *, p≤0.05; ***, p≤0.001; ****, p≤0.0001. A, E, K, M: two-way ANOVA. C: unpaired t-test. G, I: paired t-test. L: Mann-Whitney test. N: Welch’s t test.

Next, we assessed the role of the GluA1 CTD in cognitive function. We previously demonstrated that GluA1 CTD truncation does not affect spatial reference memory (Diaz-Alonso, Morishita et al. 2020). In the forced alternation Y-maze (Fig. 2B), which is used to assess spatial working memory in mice, WT and ΔCTD GluA1 male and female mice performed comparably (Fig. 2C). Then, we tested long-term spatial memory in the object location memory task (OLM, Fig. 2F). As expected from the OF results, we observed enhanced locomotion in ΔCTD GluA1 male and female mice in their first exposure to the OLM arena. To avoid its potential confounding effect, we habituated mice to the OLM arena. After 4 days, hyperlocomotion was no longer observed, indicating that ΔCTD GluA1 mice were habituated (Fig. 2D, E). Still, total locomotion during OLM training and test were significantly different between genotypes (Suppl. Fig. 2E, F), possibly driven by the introduction of novel objects in the arena. Consistent with this possibility, object exploration was also significantly greater in ΔCTD GluA1 mice during training and test (Suppl. Fig. 2G, H). Interestingly, ΔCTD GluA1 male and female mice showed superior discrimination of the displaced object compared to WT mice (Fig. 2G). We explored whether increased object exploration in ΔCTD GluA1 mice underlies their superior performance, but we found no correlation between distance travelled or object exploration time and performance in the OLM test (Suppl. Fig. 2M, N). In the novel objection recognition task (NOR, Fig. 2H), novel object discrimination was comparable between male and female ΔCTD GluA1 and WT counterparts (Fig. 2I). Neither total locomotion nor total object exploration during NOR training and test were significantly different between genotypes (Suppl. Fig. 2J-L).

### Impaired fear expression in ΔCTD GluA1 mice

Contextual fear conditioning and memory are impaired in GluA1 KO mice (Humeau, Reisel et al. 2007). Similarly, ΔCTD GluA1 mice did not exhibit freezing behavior during the conditioning phase (Fig. 2K, L). Decreased freezing was unlikely due to impaired sensory processing in ΔCTD GluA1 mice, which showed enhanced responsiveness in the hot plate test (Suppl. Fig. 2O) and higher motion indices in response to the two initial foot shocks (0.45 mA) delivered during conditioning (Suppl. Fig. 2P). Unexpectedly, ΔCTD GluA1 mice showed freezing comparable to WTs in the 24 h recall test (Fig. 2M, N), in stark contrast to GluA1 KOs, which show impaired fear expression and memory (Humeau, Reisel et al. 2007). In both WT and ΔCTD GluA1 animals, the % freezing during conditioning was not predictive of freezing during the 24 h recall test (Suppl. Fig. 2Q). These findings support that GluA1-dependent contextual memory formation does not require the CTD. Fear generalization was not affected either, supporting that context discrimination and memory function is intact in ΔCTD GluA1 mice (Suppl. Fig. 2R, S).

Next, we sought to identify the mechanism underlying the apparent discrepancy between impaired contextual fear expression (Fig. 2K, L) and intact contextual memory (Fig. 2M, N). We hypothesized that the exacerbated context novelty-driven hyperlocomotion in ΔCTD GluA1 mice masks freezing during conditioning, although it does not affect contextual memory formation. If this prediction were true, we would expect that reducing context novelty (hence decreasing hyperlocomotion) would unmask freezing during fear conditioning. We tested this by assessing contextual fear expression after a 30-min. context pre-exposure session 24 h prior to conditioning (Fig. 3A). Context pre-exposure did not affect freezing during conditioning or contextual memory in WT mice (Fig. 3B, E) but, as predicted, partially normalized freezing in ΔCTD GluA1 mice (Fig. 3B, C). As expected from previous findings (Fig. 2M, N), performance at the 24-hour recall test was indistinguishable from that of WT mice (Fig. 3D, E). Shock response was indistinguishable between ΔCTD GluA1 and WT mice in this cohort (Suppl. Fig. 3). These findings support the notion that the GluA1 CTD plays a critical regulatory role in novelty processing, but is not required for GluA1-dependent memory.

**Figure 3.**
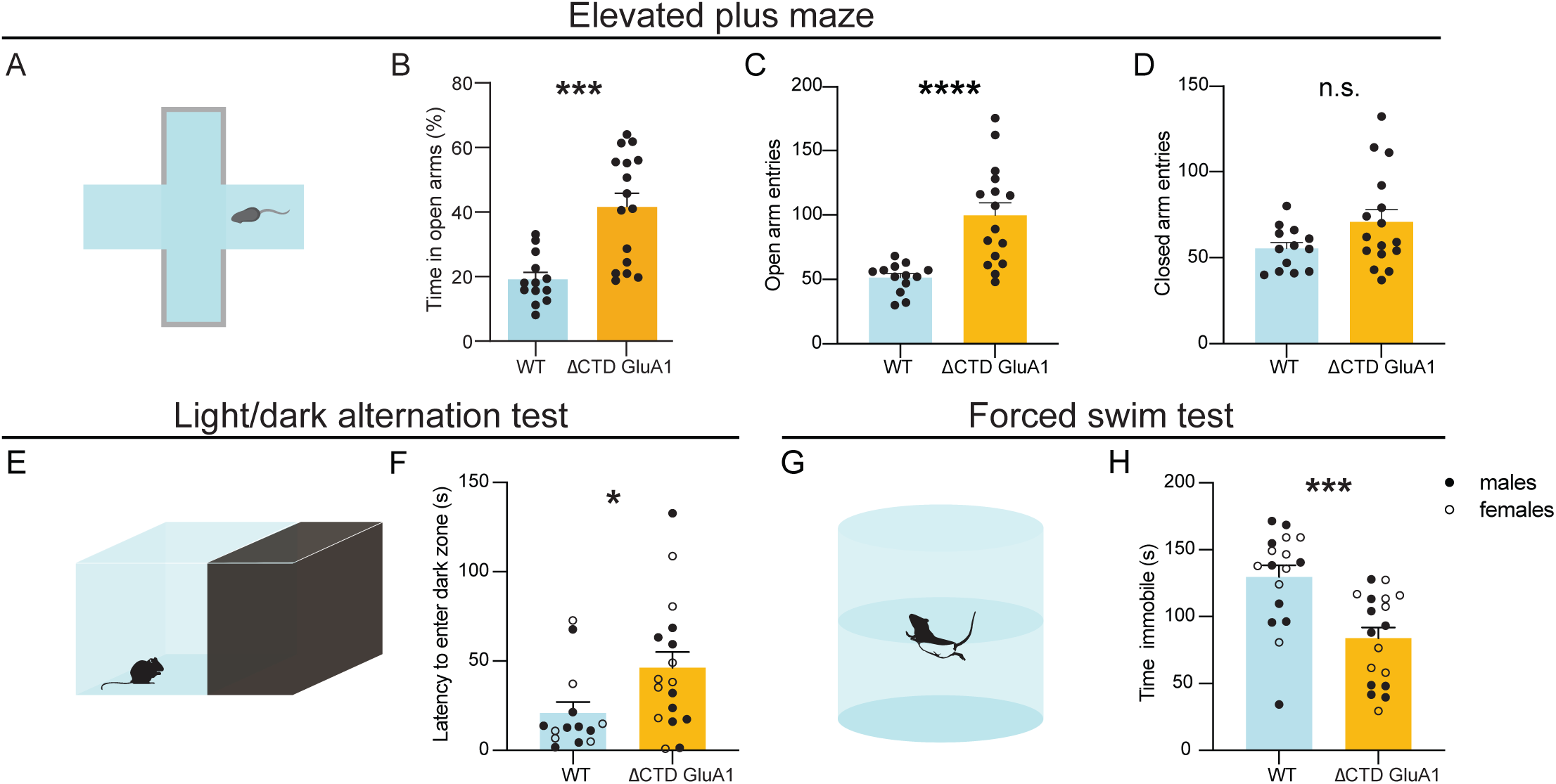
Pre-exposure to the context prior to fear conditioning partially rescues freezing behavior in ΔCTD GluA1 mice. A: Schematic of pre-exposure contextual fear conditioning paradigm. B, C: Freezing across time (B) and average freezing (C) during contextual fear conditioning for WT and ΔCTD GluA1 mice. Foot shocks are indicated with vertical red dashed lines. Horizontal dashed line indicates baseline freezing (percentage of time spent freezing during the 5 min. prior to the first shock). D, E: Freezing % across time (D) and average freezing % (E) during context recall test for WT and ΔCTD GluA1 mice. Error bars represent SEM. Empty dots represent females, filled dots represent males. n.s., not statistically different; **, p≤0.01. B, D: two-way ANOVA. C: Mann-Whitney test. E: Welch’s t-test.

### Additional schizoaffective disorder-related behavioral alterations evoked by GluA1 CTD truncation

Next, we studied whether GluA1 CTD truncation alone is sufficient to elicit other behavioral alterations relevant to schizoaffective disorders. In the elevated plus maze (EPM, Fig. 4A), ΔCTD GluA1 male mice spent a greater proportion of the time exploring the open arms (Fig. 4B) throughout the session (Suppl. Fig. 4A). Consistently, the number of open arm entries (Fig. 4C) and distance (Suppl. Fig. 4B), but not closed arm entries (Fig. 4D) and distance (Suppl. Fig. 4C) were increased in male ΔCTD GluA1 mice. Consistent with previous results (Fig. 2A, Suppl. Fig. 2D), ΔCTD GluA1 mice displayed an overall increase in total distance traveled in the EPM relative to their WT counterparts (Suppl. Fig. 4D). The observed heightened exploration of open arms in the EPM in ΔCTD GluA1 mice is reminiscent of the GluA1 KO mice phenotype (Fitzgerald, Barkus et al. 2010), albeit perhaps exacerbated. To further explore the apparently reduced anxiety in ΔCTD GluA1 mice, we applied the light/dark transition test, which can also reveal changes in anxiety-like behavior (Fig. 4E). Latency to enter the dark (safe) zone was increased in ΔCTD GluA1 male and female mice (Fig. 4F). The total time spent in each zone was not altered (Suppl. Fig. 4E). Additionally, in the forced swim test (FST, Fig. 4G), used to measure despair-like behavior in rodents, we found that ΔCTD GluA1 male and female mice spent less time immobile compared to their WT counterparts (Fig. 4H). Latency to immobility was not significantly affected (Suppl. Fig. 4F). These findings indicate that the CTD is required for GluA1-dependent novelty processing and regulates risk assessment, approach behavior and/or anxiety. Conversely, our data indicates that the CTD is not required for GluA1-dependent memory processes.

**Figure 4.**
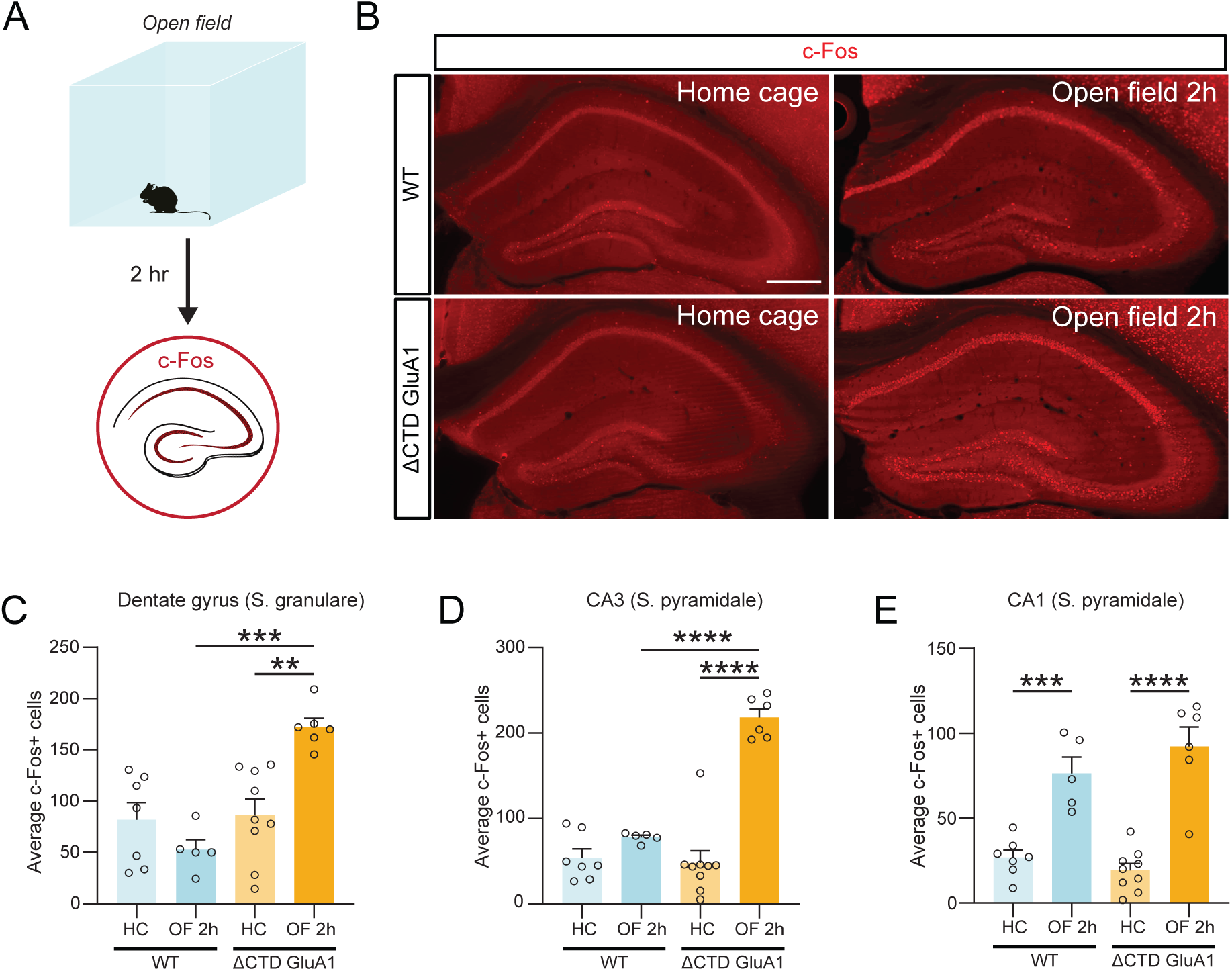
ΔCTD GluA1 mice recapitulate additional behavioral features of germline GluA1 knockout mice. A: Schematic of elevated plus maze. B-D: Mean percentage of time spent in open arms (B), total number of entries into the open arms (C) and total number of entries into the closed arms (D) for WT and ΔCTD GluA1 mice. E: Schematic of light/dark box paradigm. F: Mean latency to enter the dark compartment for WT and ΔCTD GluA1. G: Schematic of forced swim test. H: Mean time spent immobile for WT and ΔCTD GluA1 mice. Error bars represent SEM. Empty dots represent females, filled dots represent males. n.s., not statistically different; *, p≤0.05; ***, p≤0.001; ****, p≤0.0001. B, C, F: Mann-Whitney test. D, H: Welch’s t test.

### Exacerbated neuronal activity in the DG in ΔCTD GluA1 mice following exposure to a novel environment

To identify the neurobiological mechanism underlying the regulation of novelty processing by the GluA1 CTD, we sought to identify neuronal populations which respond to novelty in a GluA1 CTD-dependent fashion. To this end, we quantified c-Fos expression, a proxy for neuronal activation, two hours after exposure to a novel environment (Fig. 5A). Increased c-Fos-labelled cells were observed in various brain regions in WT male and female mice upon exposure to a novel context (Fig. 5, Suppl. Fig. 5). In dorsal hippocampus, c-Fos induction was exacerbated in putative DG GCs and field CA3 PNs in ΔCTD GluA1 male and female mice compared to WTs after OF exposure (Fig. 5B-D). c-Fos expression increased to a similar degree in WT and ΔCTD GluA1 mice in field CA1 (Fig. 5E). The similarity of these results with those previously reported in GluA1 KO mice (Procaccini, Aitta-aho et al. 2011), suggests that the CTD is critically required for GluA1-dependent regulation of hippocampal activity upon exposure to a novel context.

**Figure 5.**
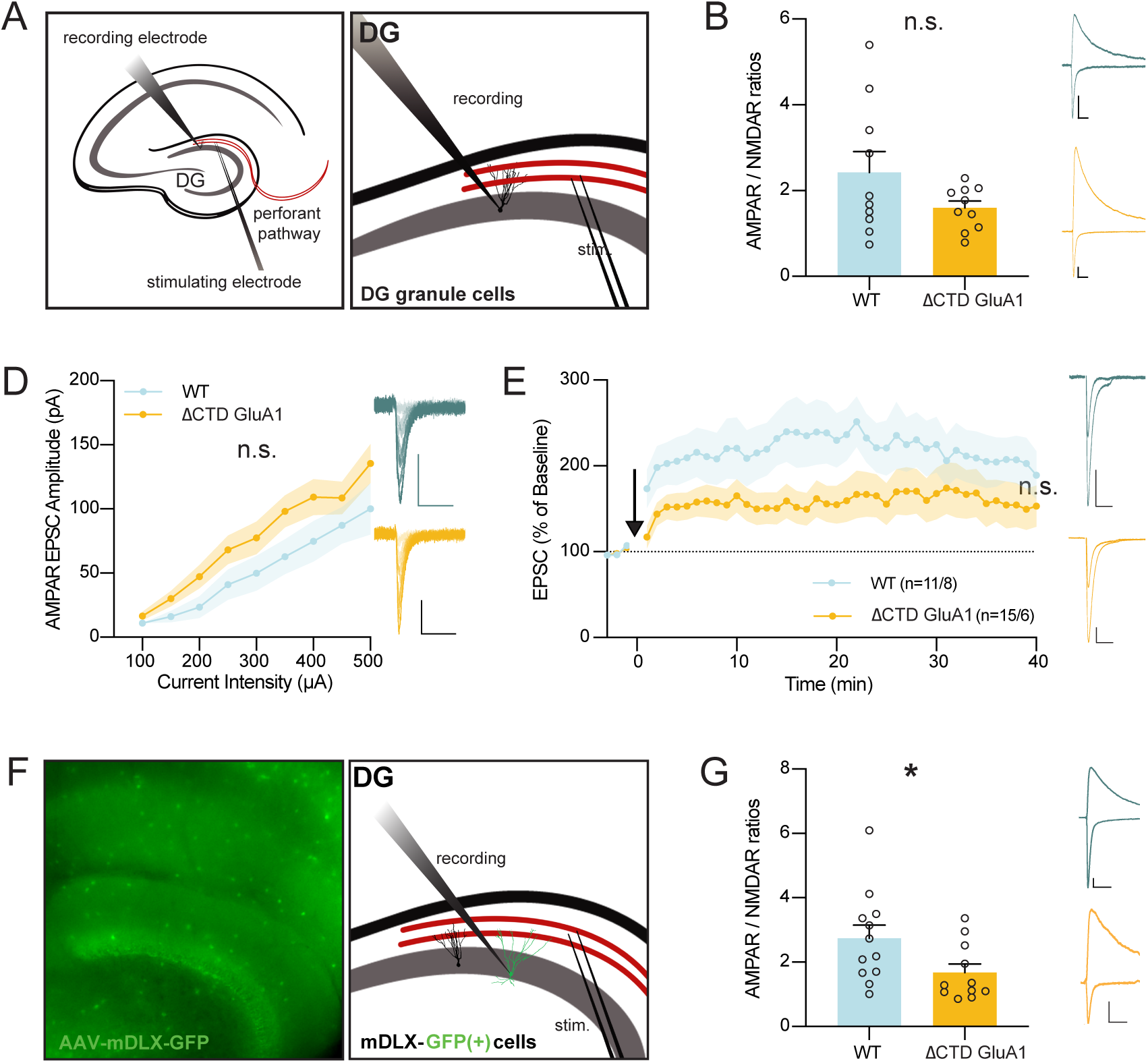
Exacerbated DG GC activation in ΔCTD GluA1 mice following open field exposure. A: Schematic of open field experiment. c-Fos expression was analyzed in several brain regions after two hours in the open field arena in WT and ΔCTD GluA1 mice. B: representative c-Fos staining (red) in WT and ΔCTD GluA1 hippocampus. C-E: Average number of c-Fos-positive cells in the dentate gyrus granule layer, CA3, and CA1, respectively. Error bars represent SEM. Empty dots represent females, filled dots represent males. Scale bar: 200 µm. **, p≤0.01; ***, p≤0.001; ****, p≤0.0001, one-way ANOVA.

### The GluA1 CTD regulates excitatory synapses onto dentate gyrus GABAergic interneurons

Excessive c-Fos expression in GCs in ΔCTD GluA1 mice can ensue as a consequence of altered synaptic transmission onto these cells. To test this possibility, we obtained whole-cell patch-clamp recordings from DG GCs using acute brain slices form ΔCTD GluA1 and WT mice (Fig. 6A) and examined excitatory synaptic transmission at perforant path (PP)➔GC synapses. We observed no significant changes in AMPAR/NMDAR ratios (Fig. 6B), indicating that AMPAR-mediated transmission is not severely affected in ΔCTD GluA1 DG GCs. Consistently, input/output AMPAR EPSC analysis showed no significant differences either (Fig. 6C), confirming that AMPAR-mediated synaptic transmission is largely intact in these cells. Then, we assessed whether the loss of the GluA1 CTD affects LTP at PP→DG GC synapses. We found a small, non-statistically significant reduction in GCs LTP in ΔCTD GluA1 mice (Fig. 6D). Altogether, these results suggest that alterations in synaptic transmission and LTP in DG GCs are unlikely to underlie the exacerbated neuronal activation observed following novel context exposure.

**Figure 6.**
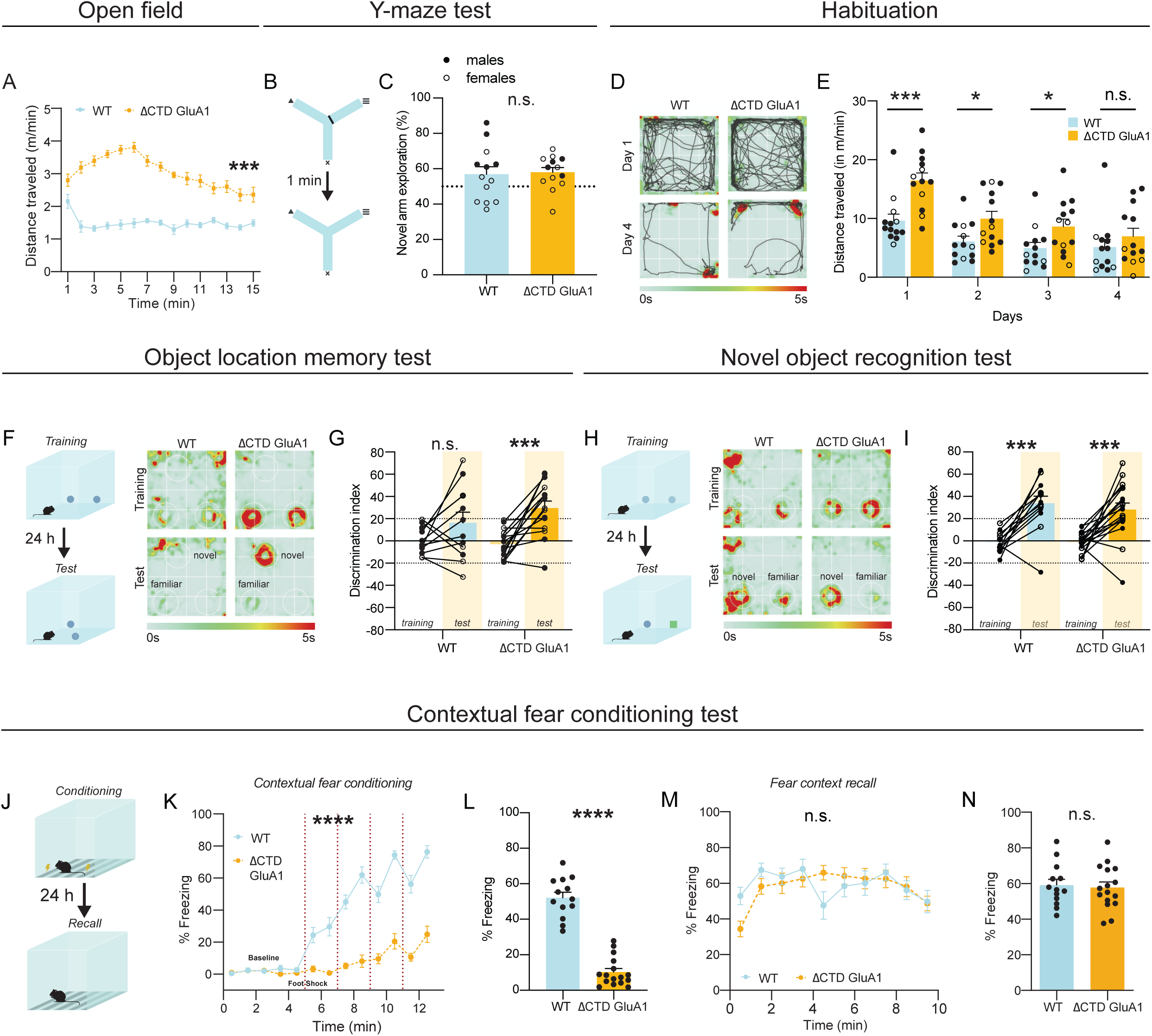
Intact excitatory synaptic transmission and LTP in DG granule cells but altered excitatory synaptic transmission in DG inhibitory INs in ΔCTD GluA1 mice. A: Whole-cell patch-clamp recording set-up for slice electrophysiology experiments in DG granule cells (GCs). B: Average paired-pulse ratio (PPR) values for evoked AMPAR EPSCs in WT and ΔCTD GluA1 GCs. Representative WT (blue) and ΔCTD GluA1 (yellow) traces are shown to the right of the plot. C: Average AMPAR/NMDAR ratios in WT and ΔCTD GluA1 GCs. D: Input-output relationship plot of AMPAR EPSCs in WT and ΔCTD GluA1 DG GCs. Representative WT (blue) and ΔCTD GluA1 (yellow) traces are shown to the right of the plot. E: AMPAR EPSC amplitude of WT and ΔCTD GluA1 DG GCs normalized to the mean AMPAR EPSC amplitude before theta-burst LTP induction (arrow). Representative WT (blue) and ΔCTD GluA1 (yellow) traces are shown to the right of the plot. n indicates number of cells induced / number of cells at the end of the experiment (min. 40). F: Whole-cell patch-clamp recording set-up for slice electrophysiology experiments in DG INs. WT and ΔCTD GluA1 mice were stereotaxically injected (AAV-mDLX-GFP) to label INs in DG. G: Mean values of AMPAR/NMDAR ratios in WT and ΔCTD GluA1 mDLX-GFP(+)-labelled INs. Representative WT (blue) and ΔCTD GluA1 (yellow) traces are shown to the right of the plot. Error bars represent SEM. Scale bars: 50pA, 20ms. n.s., not statistically different; *, p≤0.05. B-C, E, G: unpaired t-test. D: two-way ANOVA.

Local INs provide inhibitory inputs to DG GCs, thus regulating their excitability, spike timing, and lateral inhibition, and ultimately contributing to the sparse activity of DG GCs (Akgul and McBain 2016, Pelkey, Chittajallu et al. 2017, Espinoza, Guzman et al. 2018). We hypothesized that GluA1 CTD truncation might affect AMPAR-mediated excitatory synaptic transmission onto GABAergic INs in DG, thereby compromising circuit inhibition and potentially leading to the observed GCs ‘priming’. To identify inhibitory cells, we bilaterally injected an AAV-mDLX-GFP, which labels forebrain GABAergic INs, into the DG of WT and ΔCTD GluA1 littermates. After ~4 weeks of expression, GABAergic cells were labelled throughout the hippocampus in acute slices (Fig. 6E). We obtained whole-cell recordings from putative DG parvalbumin (PV)+ basket cells, identified by their morphology and localization of the soma within SG. We found a significant reduction in AMPAR/NMDAR ratios in these cells (Fig. 6F), indicating that the loss of the GluA1 CTD affects synaptic transmission in DG GABAergic INs, in contrast to the intact synaptic transmission observed onto GCs. The specific reduction of excitatory synaptic drive onto DG GABAergic cells explains, at least in part, the exacerbated DG responsiveness to novelty and subsequent behavioral alterations observed in ΔCTD GluA1 mice.

## Discussion

GluA1-deficient mice exhibit deficits in synaptic plasticity and behavioral alterations, such as selective deficits in short-term habituation and exacerbated novelty-induced locomotor hyperactivity, reminiscent of some of the features of schizoaffective disorders and neurodevelopmental conditions including attention-deficit/hyperactivity disorder (Fitzgerald, Barkus et al. 2010, Barkus, Feyder et al. 2012, Barkus, Sanderson et al. 2014). Consistently, mutations in the *GRIA1* gene, which encodes GLUA1, may increase risk of schizophrenia in humans (Coyle 2006, Ripke, O’Dushlaine et al. 2013, Schizophrenia Working Group of the Psychiatric Genomics 2014, Ismail, Zachariassen et al. 2022, Yonezawa, Tani et al. 2022).

What makes GluA1 unique among AMPAR subunits? The GluA1 CTD is the most sequence-diverse area of the receptor and has therefore drawn considerable attention for decades. Despite the interest, its role, especially at synapses outside of hippocampal field CA1, is largely unexplored. In this study, we used constitutive GluA1 CTD-truncated mice to explore crucial aspects of how the CTD affects GluA1’s localization and function at the biochemical, cellular and behavioral level. We found that the GluA1 CTD regulates AMPAR subunit protein levels, intracellular trafficking and synaptic transmission onto inhibitory, but not excitatory neurons in the DG, ultimately affecting GC excitability and spatial novelty processing. We found no evidence of memory impairments upon loss of the GluA1 CTD, and in fact we observed enhanced performance in OLM. Altered performance in the FST, EPM and light/dark alternation tests suggest additional regulation of affective processes by the GluA1 CTD.

In a previous study we did not observe qualitative changes in AMPAR subunit expression in ΔCTD GluA1 mice (Diaz-Alonso, Morishita et al. 2020). However, more detailed analysis in this study revealed that GluA1 subunit levels and subcellular distribution are, in fact, affected by the loss of the GluA1 CTD. We also found that the CTD influences intracellular GluA1 trafficking, consistent with previous reports highlighting the importance of GluA1 CTD interactions with 4.1N and SAP97 in intracellular AMPAR trafficking (Shen, Liang et al. 2000, Sans, Racca et al. 2001, Bonnet, Charpentier et al. 2023). Interestingly, despite reduced GluA1 levels and altered intracellular trafficking, we found that both GluA1’s abundance at synaptosomes and its colocalization with PSD-95 were not significantly affected by truncation of the CTD. These findings suggest that, despite reduced soma→dendrite trafficking, synaptic AMPAR docking is not significantly affected by the truncation of the GluA1 CTD. This is consistent with the normal AMPAergic transmission in ΔCTD GluA1-expressing CA1 PNs (Granger, Shi et al. 2013, Diaz-Alonso, Morishita et al. 2020, Watson, Pinggera et al. 2021) and DG GCs (present study).

GluA2 protein levels were dramatically increased in ΔCTD GluA1 mice, in stark contrast with the unaltered or even reduced GluA2 levels reported in GluA1 KO mice (Zamanillo, Sprengel et al. 1999, Jensen, Kaiser et al. 2003). Furthermore, GluA2, but not GluA3 subunits, also appeared enriched in the soma in ΔCTD GluA1 mice, suggesting that GluA2 can form stable heteromeric receptors with ΔCTD GluA1 and that the GluA1 CTD exerts a significant influence in intracellular trafficking of GluA1/A2 AMPARs. Altogether, these findings support the notion that the GluA1 subunit, both via its ATD (Diaz-Alonso, Sun et al. 2017) and its CTD (present study), dominate heteromeric AMPAR trafficking. Together with the normal levels and localization observed for GluA3, and the unaltered GluA2/A3 colocalization in ΔCTD GluA1 hippocampi, these findings suggest that CTD-lacking GluA1 partakes in synaptic transmission similarly to WT GluA1, and that the normal synaptic transmission and plasticity observed at CA1 PNs and DG GCs are not a result of a replacement of GluA1-containing AMPARs by GluA2/A3 heteromers.

The mechanisms regulating AMPAR trafficking and synaptic complement are poorly understood outside of hippocampal field CA1, despite the prevalence of AMPAR-mediated synaptic transmission throughout the CNS. Here we found that DG GCs are “primed” in ΔCTD GluA1 mice, and become excessively active following spatial novelty exposure, presumably contributing to hyperlocomotion. A recent study offered a plausible explanation for GC overactivity in ΔCTD GluA1 mice, showing that AMPAR EPSCs are enhanced in GCs overexpressing CTD-lacking GluA1, which escapes SAP97-mediated retention at perisynaptic sites (Kay, Tsan et al. 2022). In this study, we did not find increased AMPAR EPSCs in ΔCTD GluA1 mice, possibly because of the different approach (constitutive GluA1 CTD truncation *vs* acute overexpression of CTD-truncated GluA1) or species (mouse *vs* rat) employed in the two studies. Instead, we found an alternative possibility: AMPAR EPSCs on DG inhibitory INs are significantly smaller in ΔCTD GluA1 mice, which conceivably leads to decreased inhibition onto DG GCs and may thereby render DG GCs prone to overactivation by excitatory inputs, especially those conveying novelty. These findings are consistent with a previous report showing that chemogenetic hippocampal inhibition normalized novelty-induced locomotion in GluA1 KO mice (Aitta-Aho, Maksimovic et al. 2019). Our results suggest that, while altered AMPAR subunit levels and intracellular trafficking affect various neuron types in ΔCTD GluA1 mice, certain AMPAR subunit compositions, such as the GluA1/GluA4 heteromers that dominate in fast-spiking PV+ INs, are particularly sensitive to the truncation of the GluA1 CTD. Meanwhile, excitatory neurons may more easily compensate the truncation of the GluA1 CTD. The increased levels of GluA4, whose expression is essentially restricted in the forebrain to PV+ INs, is additional support for their specific vulnerability in the ΔCTD GluA1 DG. Alternatively, it may hint a compensatory mechanism involving this cell population.

PV+ INs dysfunction can contribute to the pathophysiology of schizophrenia (Lisman, Coyle et al. 2008, Curley and Lewis 2012, Marin 2012, Ruden, Dugan et al. 2021). Altered AMPAR function in PV+ INs can significantly affect their output and function, as exemplified in PV+ IN-specific GluA1 KO mice, which show impaired short-term habituation (Fuchs, Zivkovic et al. 2007), and excitation/inhibition imbalance reminiscent of that found in patients with schizophrenia (Chen-Engerer, Jaeger et al. 2022). Other manipulations such as the deletion of Erbb4 in PV+ INs, which lead to a reduction in AMPAR content in excitatory synapses onto PV+INs, also result in schizophrenia-related phenotypes (Del Pino, Garcia-Frigola et al. 2013). The important role of the GluA1 CTD supporting excitatory synapses onto putative PV+ INs unveiled in this study expands our understanding of the mechanisms underlying cell type-specific AMPAR transmission, disruptions of which potentially contribute to altered synaptic transmission in schizoaffective disorders.

Our study discriminates between CTD-dependent and independent GluA1 cognitive processes: on one hand, we demonstrate that spatial working memory, object recognition memory and long-term contextual fear memory – all of which are impaired in GluA1 KO mice (Reisel, Bannerman et al. 2002, Humeau, Reisel et al. 2007, Sanderson, Good et al. 2009), are not affected by the loss of the GluA1 CTD. Remarkably, OLM is enabled after subthreshold training. On the other hand, we find that GluA1 CTD truncation alone is sufficient to reproduce aberrant salience, short-term habituation and general response to novelty. The normalization of fear expression during contextual fear conditioning by context pre-exposure suggests that disrupted fear response in ΔCTD GluA1 mice is secondary to altered novelty processing. Altogether, our findings clearly demonstrate that GluA1-dependent regulation of novelty processing necessitates the CTD.

GluA1 KO mice are considered a valuable tool to study altered synaptic function in schizophrenia (Fitzgerald, Barkus et al. 2010, Barkus, Feyder et al. 2012, Bygrave, Jahans-Price et al. 2019). Here we found that GluA1 CTD truncation alone recapitulated the schizoaffective-relevant behaviors present in GluA1 KO mice. Specifically, the increase in approach behavior in the elevated plus maze, light/dark transition and forced swim tests can be interpreted as reduced anxiety / depression, but may also reflect increased novelty-seeking or risk-taking, recapitulating and even exacerbating some of the symptoms of schizophrenia and ADHD previously observed in constitutive GluA1 KOs. Similar to genetic deletion of GluA1, the behavioral consequences of GluA1 CTD truncation are complex, and a complete, accurate interpretation will require additional studies.

In summary, this study provides a comprehensive characterization of the GluA1 CTD roles in AMPAR subunit levels, intracellular trafficking, cell type-specific synaptic transmission and GluA1-dependent affective and memory processes. Our study identifies the GluA1 CTD as a crucial element in the AMPAR complex that regulates the strength of excitatory synapses onto inhibitory INs, and suggests that ΔCTD GluA1 mice may be valuable to study features of schizoaffective and other psychiatric disorders.

## Acknowledgments

We would like to thank Dr. Roger Nicoll for supporting initial experiments in his laboratory. Dr. Mulatwa T. Haile and Dr. Lulu Y. Chen for guidance and equipment used in behavior assessments, and the Diaz-Alonso lab members for fruitful discussions. This work is supported by grants K99/R00 MH118425, Whitehall Foundation, Brain and Behavior Research Foundation and UCI start-up funds to J.D.-A. and AG076835 to M.A.W. G.S. is supported a the T32 Training Program in Epilepsy Research (T32NS045540). C.A.C. is supported by an HHMI Gilliam’s Fellowship. A.M. is supported by the NIH-NIGMS Maximizing Access to Research Careers T34 (#GM136489). M.A.S. is supported by a Eugene Cota-Robles fellowship and the Howard Schneiderman T32 Training Program in Learning and Memory (#T32MH119049). V.A.V. is supported by the NRSA DA059982 fellowship. The Optical Biology Core Facility of the Developmental Biology Center is supported by grants CA-62203 and GM-076516.

## Author contributions

G.S. performed and analyzed electrophysiology experiments; A.V.K. performed and analyzed biochemistry experiments; G.S., A.V.K and M.A.S. performed and analyzed histology experiments; G.S., A.V.K., C.A.C., A.M., V.A.V., I.L., J.S. and M.A.W. performed and analyzed behavior experiments; G.S and J.D.-A. drafted, and all authors edited the manuscript. J.D.-A. coordinated the study.

## Figure legends

**Suppl. Figure 1. Analysis of excitatory synapse density in CA1 and DG in WT and ΔCTD GluA1 mice.**

A-B: Average density of GluA1 and PSD-95 positive puncta in CA1 SR. C-D: Average density of GluA1 and PSD-95 positive puncta in DG ML. E-F: Average density of GluA2 and GluA3 positive puncta in CA1 SR. G-H: Average density of GluA2 and GluA3 positive puncta in DG ML. Error bars represent SEM. n.s., non-statistically significant; *, p≤0.05, unpaired t-test.

**Suppl. Figure 2. Control behavioral assessments in WT and ΔCTD GluA1 mice (related to Fig. 2).**

A-C: Average thigmotaxis (A), fine movements (B), and rearings (C) of WT and ΔCTD GluA1 male mice during an open field test. D: Total distance travelled of WT, heterozygous, and homozygous ΔCTD GluA1 female and male mice in the open field test. E-F: Mean distance traveled during training (E) and test (F) for WT and CTD GluA1 mice in the OLM task. G-H: Mean object exploration time during training (G) and test (H) for WT and CTD GluA1mice in the OLM task. I-J: Mean distance traveled during training (I) and test (J) for WT and CTD GluA1 mice in NOR task. K-L: Mean object exploration time during training (K) and test (L) for WT and CTD GluA1 mice in NOR task. M: Linear regression of total distance traveled (meters) and discrimination index during OLM test day. N: Linear regression of total object exploration (seconds) and discrimination index during OLM test day. O: Average hind paw withdrawal latency of WT and CTD GluA1 mice in the hot plate test. P: Average motion index of WT and ΔCTD GluA1 mice during contextual fear conditioning (arbitrary units). Q: Linear regression of percentage of freezing during conditioning (10 min) and recall (8 min). R: Average percentage of freezing measured across time (minutes) during the fear generalization test for WT and ΔCTD GluA1 mice. S: Average percentage of freezing during the fear generalization test for WT and ΔCTD GluA1 mice. Error bars represent SEM. Empty dots represent females, filled dots represent males. n.s. not statistically different, *p≤0.05, ***p≤0.001. A, B, H, O, S: Welch’s t test. C, I, J, K: Mann-Whitney test. D: one-way ANOVA. E-G, L: unpaired t-test. M, N, Q: Linear regression. P: multiple t design. R: two-way ANOVA.

**Suppl. Figure 3. Shock reactivity of WT and ΔCTD GluA1 mice in contextual fear conditioning paradigm.**

Average motion index of WT and ΔCTD GluA1 mice during contextual fear conditioning (arbitrary units). Error bars represent SEM. Empty dots represent females, filled dots represent males. Non-statistically significant differences, two-way ANOVA.

**Suppl. Figure 4. Control behavioral assessments in WT and ΔCTD GluA1 mice (related to Fig. 4).**

A: Average time spent in the open arms across time during the elevated plus maze for WT and ΔCTD GluA1 mice. B: Ratio of distance traveled in open arms relative to total distance traveled during the elevated plus maze for WT and ΔCTD GluA1 mice. C: Average distance traveled in the closed arms during the elevated plus maze for WT and ΔCTD GluA1 mice. D: Average total distance traveled during the elevated plus maze for WT and ΔCTD GluA1 mice. E: Average time spent in the light zone of the light/dark alternation test for WT and ΔCTD GluA1 mice. F: Average latency to immobility during the forced swim test for WT and ΔCTD GluA1 mice. Error bars represent SEM. Empty dots represent females, filled dots represent males. n.s., non-statistically significant; **, p≤0.01, ****, p≤0.0001, A: two-way ANOVA. B-D: Welch’s t-test. E: Mann-Whitney test. F: unpaired t-test.

**Suppl Figure 5. c-Fos analysis in various brain regions of WT and ΔCTD GluA1 mice following open field exposure.**

A: Schematic of experimental timeline. B: c-Fos staining (red) of representative WT and ΔCTD GluA1 mouse brains showing habenula, somatosensory cortex, subthalamic nucleus, and amygdala. C: c-Fos staining (red) of representative WT and ΔCTD GluA1 mouse brains showing prefrontal cortex. D: c-Fos staining (red) of representative WT and ΔCTD GluA1 mouse brains showing motor cortex, striatum, and nucleus accumbens. Error bars represent SEM. Error bars represent SEM. Empty dots represent females, filled dots represent males. n.s., not statistically different; *, p≤0.05; **, p≤0.01; ***, p≤0.001; ****, p≤0.0001, unpaired t-test, one-way ANOVA.

