## Supplementary figures and images for "The GluA1 cytoplasmic tail regulates intracellular AMPA receptor trafficking and synaptic transmission onto dentate gyrus GABAergic interneurons, gating response to novelty"

### Supplemental Figure 1

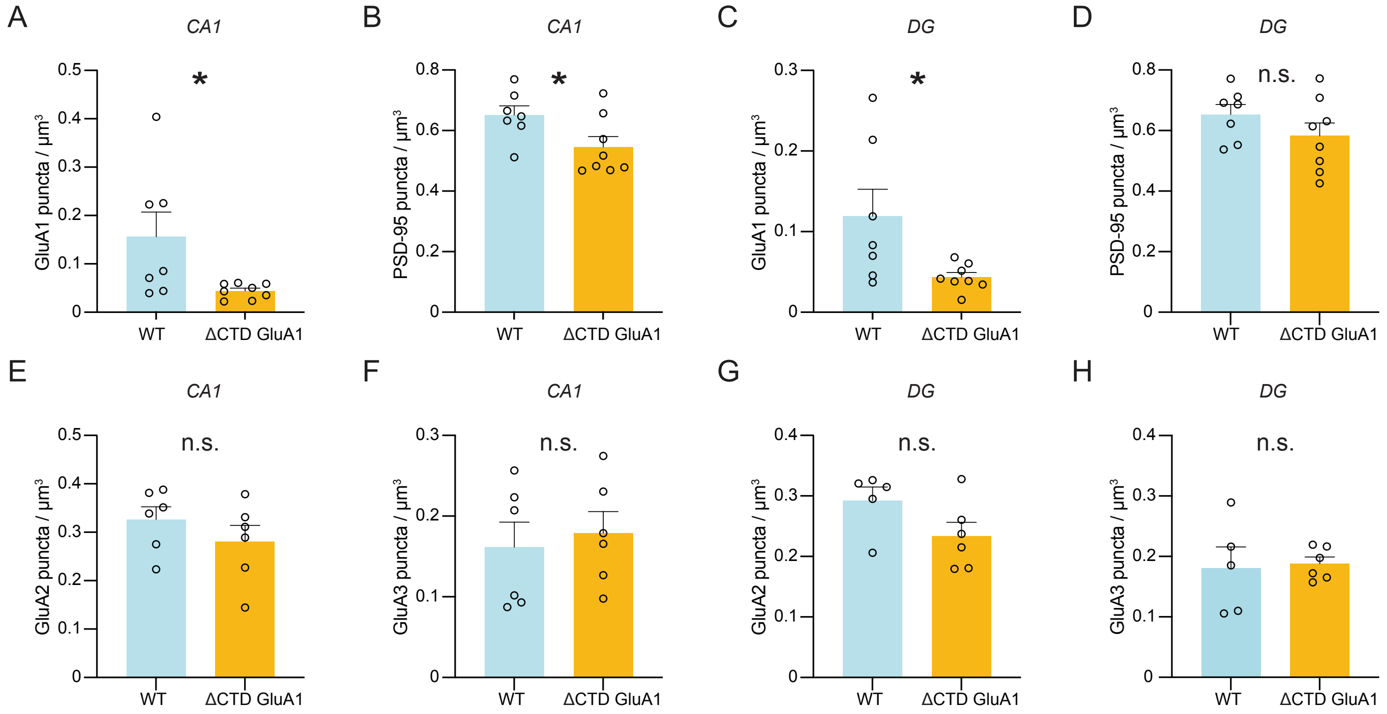

### Supplemental Figure 2

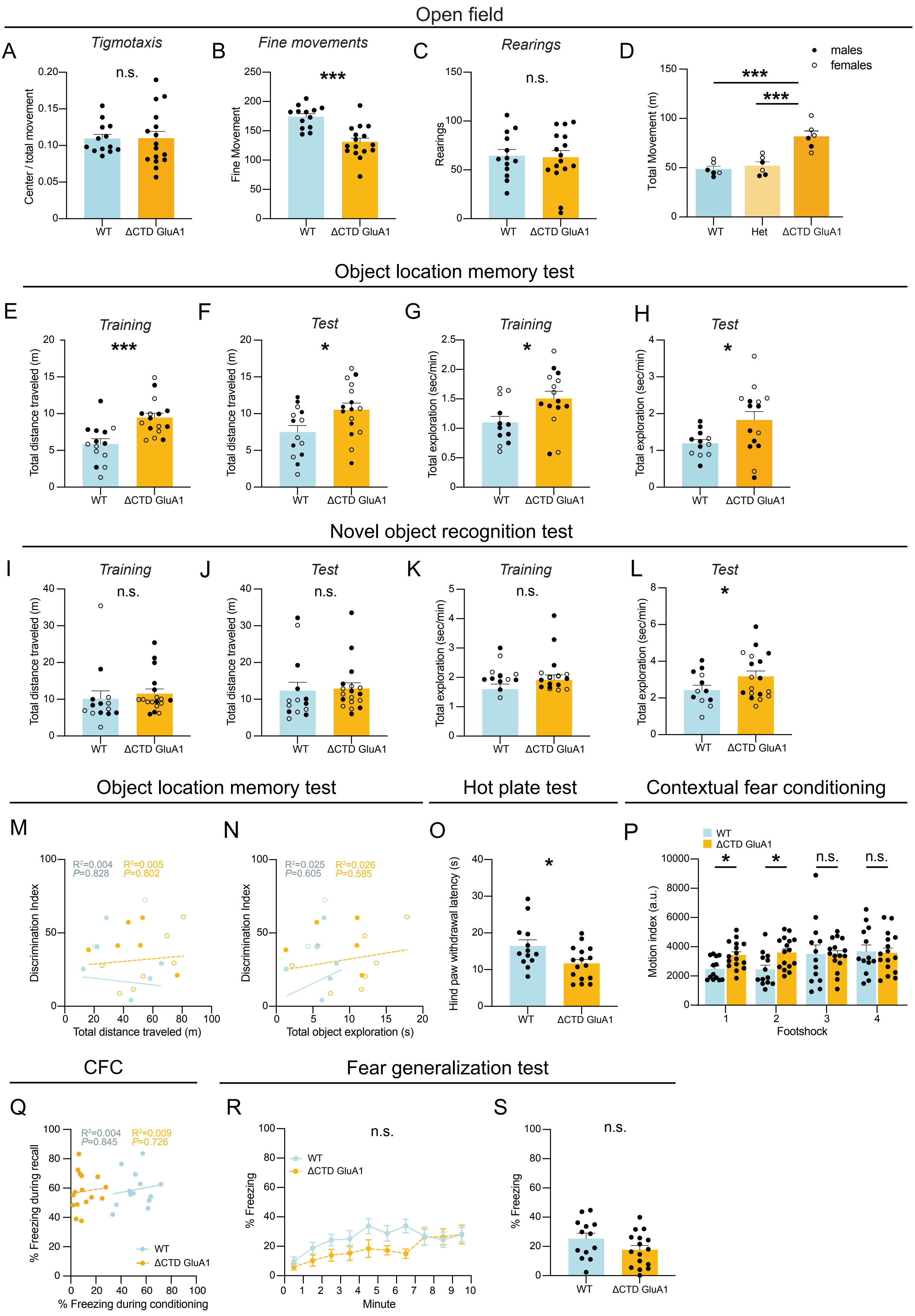

### Supplemental Figure 3

*Contextual Fear Conditioning Test - Pre-exposure*

A

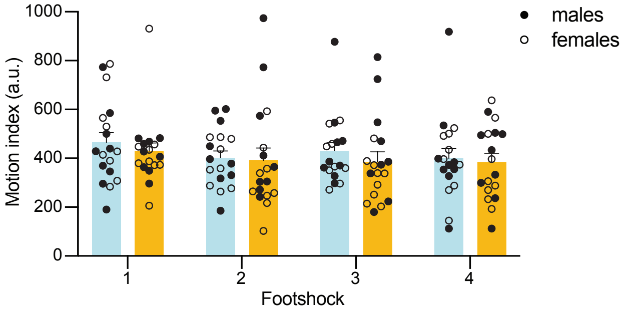

### Supplemental Figure 4

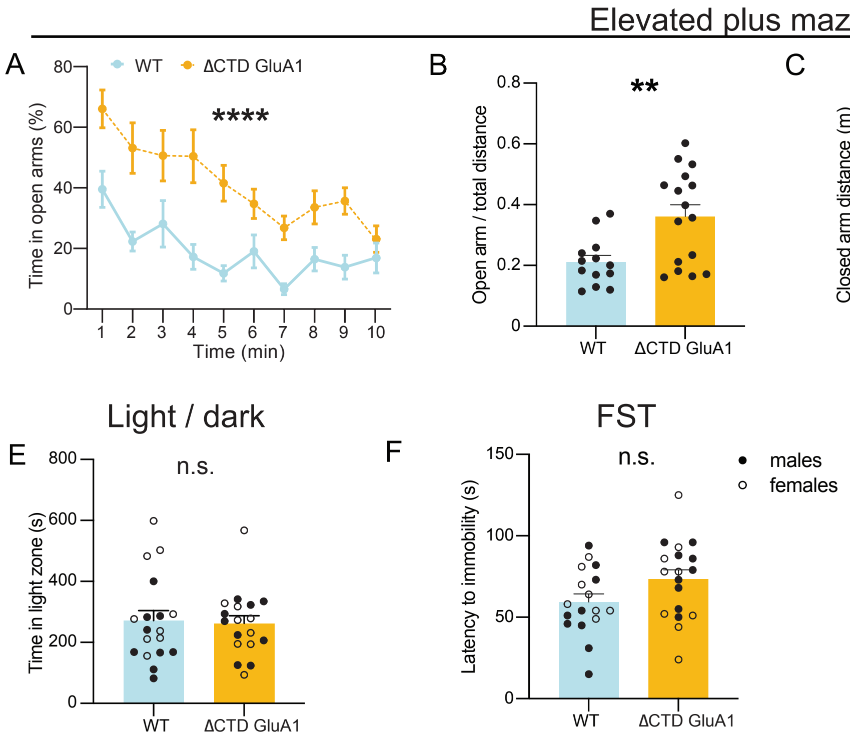

### Supplemental Figure 5

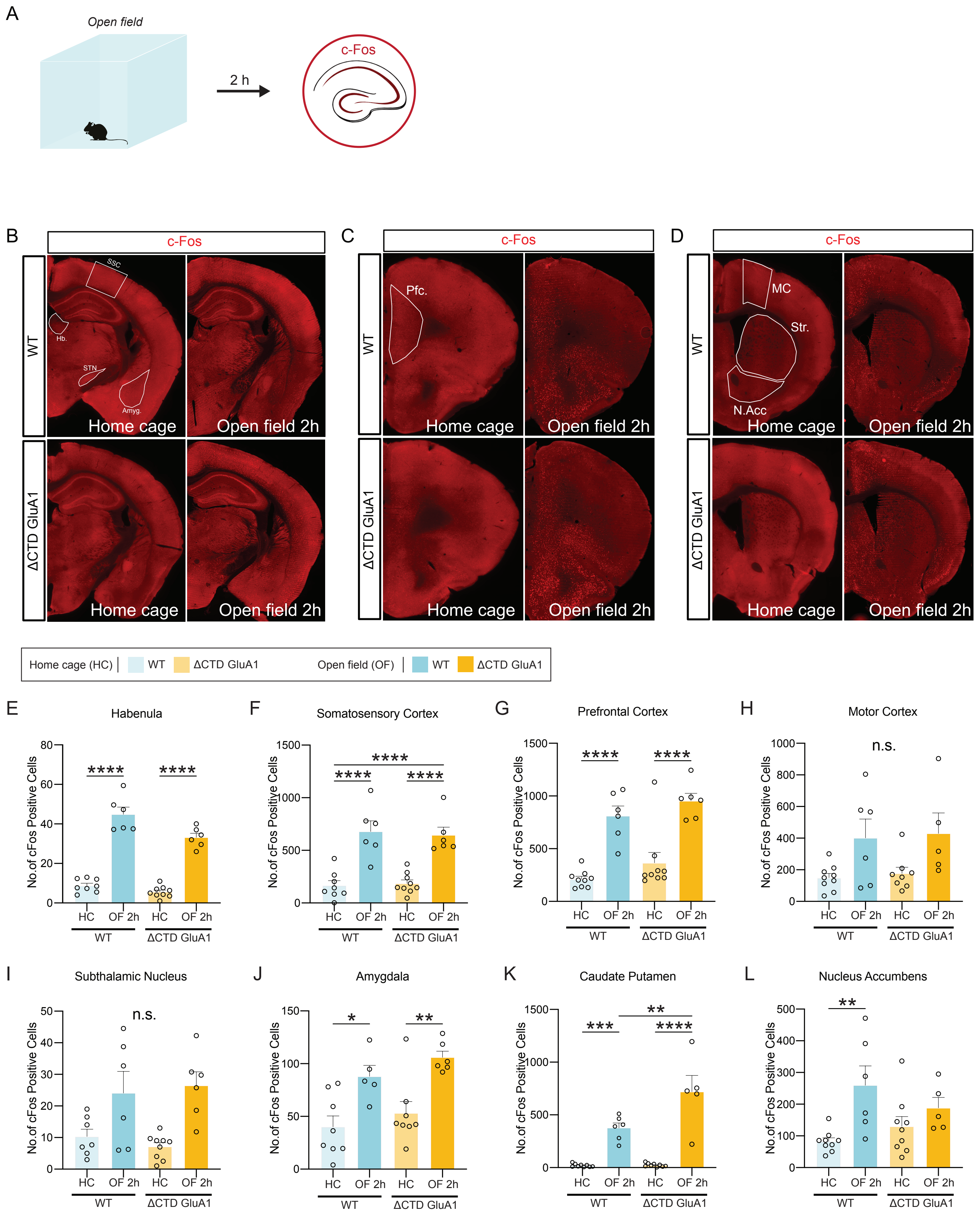
